# Reinforcement Learning Based Approach for Ligand Pose Prediction

**DOI:** 10.1101/2021.04.10.438538

**Authors:** Justin Jose, Kritika Gupta, Ujjaini Alam, Nidhi Jatana, Pooja Arora

**Affiliations:** Engineering for Research, ThoughtWorks Technologies, Pune, 411006, India; CSIR-Institute of Genomics and Integrative Biology, New Delhi, 110025, India

**Keywords:** protein-ligand docking, reinforcement learning, DDPG, GraphCNN, physico-chemical properties, drug discovery, optimization

## Abstract

Identification of the potential binding site and the correct ligand pose are two crucial steps among the various steps in protein ligand interaction for a novel or known target. Currently most of the deep learning methods work on protein ligand pocket datasets for various predictions. In this study, we propose a reinforcement learning (RL) based method for predicting the optimized ligand pose where the RL agent also identifies the binding site based on its training. In order to apply various reinforcement learning techniques, we suggest a novel approach to represent the protein ligand complex using graph CNN which would help utilize both atomic and spatial features. To the best of our knowledge, this is the first time an RL based approach has been put forward for predicting optimized ligand pose.

## I. Introduction

The drug discovery process is long, challenging and expensive. It is the first step in the identification of new candidate molecules for treating human disease. According to research reports, average cost and time to bring one drug molecule to market is approximately $2 billion and 10-12 years respectively [1]. Another study reveals that this cost has doubled since 2010. Completion of human genome has helped in figuring out a number of targets [2], which has escalated the need to identify the corresponding leads for target proteins. The expense incurred due to the time scale is also augmented by the compute expense, as lead identification and the study of protein-ligand interaction is one of the most compute expensive processes.

The COVID-19 pandemic has unveiled a pressing need to develop effective drugs rapidly [3]. It has indicated a need for expediting efficient computational methods for drug discovery. To address this need, the Government of India launched a drug discovery hackathon (DDH)^1^ in July 2020 to develop computational methods for COVID-19 drug discovery, and to extrapolate these algorithms to other generic drug discovery challenges. In this work, we explore one of these problems statements which requires the application of reinforcement learning algorithms to the prediction of ligand pose during protein ligand interaction.

Computational methods for identification of the right ligand pose have been explored for a while. Structure based virtual screening methods, such as molecular docking, have been widely used for identification of the lead molecule for virtual screening [4]. Until the last few years, molecular docking was the only method for protein ligand interaction identification. In this, most of the docking tools focus towards calculation of binding energy or binding affinity, where the ligands and their poses are then ranked according to their score. A common challenge is that often the ligand pose chosen by these methods need to be manually checked, as a good score may or may not lead to the correct biological interaction.

### A. Literature Survey

The literature on molecular docking presents various approaches. Majority of these docking approaches follow an algorithmic approach to identify docking poses for the small molecules, taking into account the molecular interaction between the protein and ligand molecules [4], [5]. The goodness of the approximated docking function is validated with an associated scoring function [6].

Given the various challenges faced in molecular docking, over the last couple of years there has been a surge in the usage of AI/ML methods for protein ligand interaction. Most machine learning based approaches designed to predict the protein-ligand binding affinity try to approximate functions which can reliably be used to score the goodness of the poses generated using empirical methods [7]–[10]. In all these methods, the quality of the predicted pose is heavily influenced by the underlying scoring function, which in itself forms a separate field of research.

Reinforcement learning, which is a known technique in the field of optimization, has also made some recent progress in the protein structure world. The following section briefly explains the current role of reinforcement learning in drug design and the kind of molecular representations required to apply various reinforcement learning methods.

#### 1) Role of Reinforcement Learning in Drug Design

In the existing literature, reinforcement learning has been predominantly used as a supplementary aid in generative drug design. Some of the issues in generative drug design algorithms are alleviated with the help of targeted optimization using fine-tuned, task-specific reward functions [11]. In a study by Olivecrona et al. [12], the reinforcement learning approach was utilized to generate molecules which reflect the intent of a scoring function. This was achieved by training the agent with the estimation of molecule quality under a pre-trained network in chemical space and with a scoring function to guide the model in producing output examples which are still representative of the original chemical domain. It was observed that the agent could produce structurally similar analogues, even when all such compounds were removed from the training set.

Reinforcement learning requires featurized inputs for state representation. Generally, the input state can be represented as linear vectors, or feature vectors extracted using a convolution network. In case of molecular structures, a 3D CNN can be used [8] to better represent the spatial features of the molecule. Computationally, 3D CNNs are expensive, hence a faster alternative needs to be identified, which can be used to extract feature vectors from molecule representations.

#### 2) Molecule representation as Graph Convolution Networks

Recently, graph convolutions on molecules have gained popularity due to their ability to process molecules of varying size into learnable feature vectors (fingerprints) of fixed length [13]–[16]. They work on molecules represented as 2D graphs– atoms as nodes embedded with atomic features, and bonds as edges. Just like convolutions on images process pixels and their receptive fields to generate an output, convolutions on graphs process nodes and their neighbors (i.e., atoms and their bonded atoms) to generate an output that incorporates the atom-atom interactions.

A general architecture for convolutions on molecular graph inputs is defined in earlier works [13], for which open-source implementations in Tensorflow [17] exist in the form of three layers, namely, Graph Convolution, Graph Pooling, and Graph Gathering [14].

## II. Approach

In this study, we put forward a method that employs reinforcement learning for the identification of optimized ligand pose. The overall approach, as shown in figure 1, comprises of the following steps.

- Molecular representation of protein-ligand complex: This involves identifying a suitable form to represent the protein-ligand complex, which would aid in applying reinforcement learning techniques to it.
- Prediction of optimized ligand pose: This avails the protein and ligand atomic and spatial features, and their inter molecular interactions at the binding site, to identify optimal ligand pose. In our formulation, the reinforcement learning agent is responsible for identifying both the binding site and the optimized ligand pose. We propose to train the reinforcement learning model based on existing protein-ligand complexes.

**Fig. 1.**
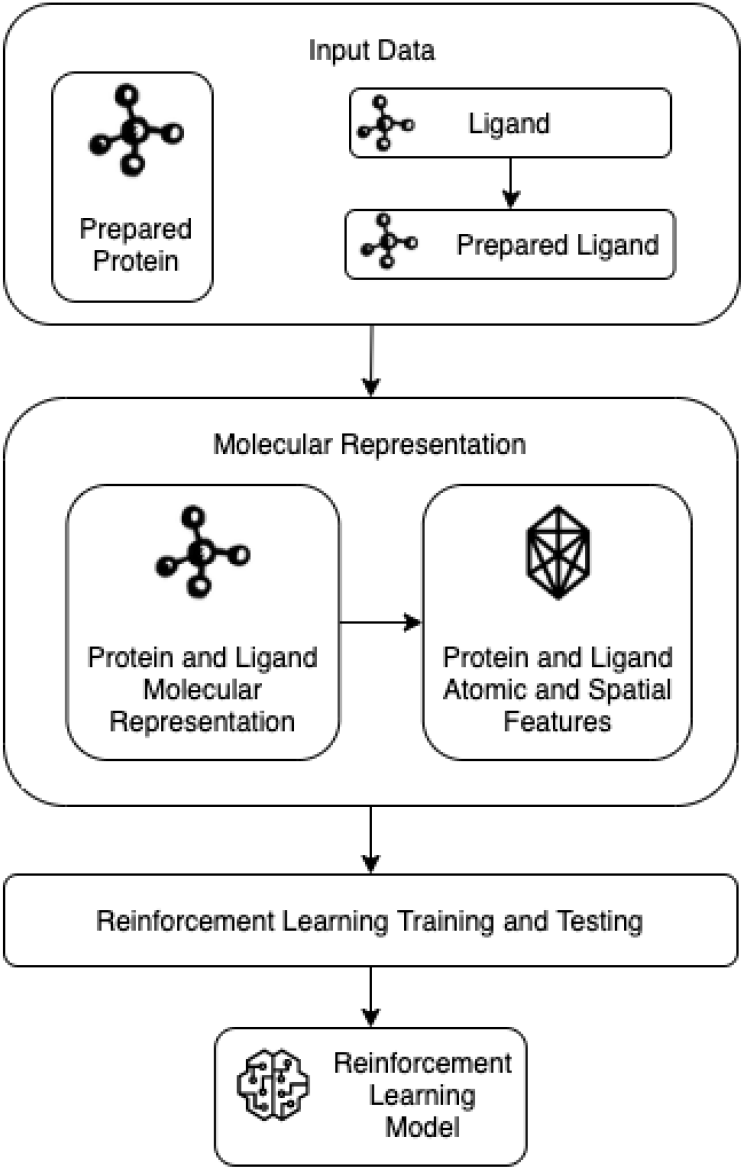
Overall workflow including: 1. Getting protein ligand complex dataset 2. Representing the dataset in 4D space 3. RL training and 4. RL model.

The hypothesis presented is that training on protein-ligand complexes with known binding pose will help the reinforcement learning algorithm to approximate the underlying molecular interactions by making use of the input atomic and spatial features presented as molecule fingerprint. This would lead to an optimized pose in the desired binding site of the protein of interest, based on the atomic, spatial and molecular features.

### A. Protein-Ligand Docking as an optimization function

Protein-ligand docking can be viewed as a combinatorial optimization problem [18] which involves searching and selection sub-stages. Deep learning approaches are function approximators [19] which are capable of identifying underlying feature interactions provided as part of the input features. Reinforcement learning approximates an optimization function in the form of optimal policy [20], which helps in maximizing the total reward. Deep reinforcement learning strategies combine deep learning function approximation with reinforcement learning’s optimization capabilities. In the case of protein-ligand docking, the reinforcement learning algorithm approximates the pose prediction function. The optimal policy learned by the algorithm generates the ligand pose which can improve the protein-ligand interaction at binding site, thus implicitly combining the searching and selection process, which earlier existed as separate, explicit processes.

The ligand pose generated as part of the reinforcement learning is a spatial vector representing the relative position and alignment of the ligand with respect to the protein. This spatial vector is continuous in nature, hence the chosen reinforcement learning algorithm needs to operate in the continuous state-space. Deep Deterministic Policy Gradient (DDPG) [21] is a continuous state and action space reinforcement learning algorithm developed using DPG [22], which may be appropriate for this purpose.

## III. Materials and Methods

In this section, we elaborate on the dataset used to understand the problem, and the network design, which includes data representation and RL process.

### A. Collating Protein-Ligand Data

The dataset consists of 13 viral protein ligand complexes selected from PDBbind [23] and RCSB PDB [24]. As expected in the problem statement from DDH, the dataset includes viral protein-ligand complexes which are cysteine protease, serine protease, helicase and RNA dependent RNA polymerase. The complexes were prepared using AutoDockTools (ADT) [25], [26]. To study the viral protein ligand interactions required for training set, we chose papain-like protease (PLpro) and 3C-like main protease (3CLPro) where a set of 15 ligands each for papain-like protease (PLpro) and 3C-like main protease (3CLPro) were mined from the literature [27]–[30] and the 2D structures were downloaded from Pubchem [31]. Open babel was used to generate the 3D structures of all the ligands [32]. The ligand and protein preparation was carried out using AutoDockTools (ADT). Autodock vina was used to carry out docking of ligands with the viral proteins [33] to understand the binding site and interactions at the binding site.

Based on this initial analysis and previous work, the following primary features can be selected to represent the protein ligand interaction: 3D coordinates (along x, y and z axes), atomic number, hybridization, heavy valency, hetero valency, partial charge, hydrophobicity, aromaticity, hydrogen acceptor, hydrogen donor, and ring. These can be extracted using the OpenBabel tool. Eventually more specific features can be added based on the output of initial trials.

### B. Network Design

The overall network for the reinforcement learning based protein-ligand docking comprises of a GraphCNN layer responsible for representing atomic and molecular properties as a feature vector, and an optimization function approximating the docking function, which is derived using the reinforcement learning layer.

#### 1) Data Representation

For the representation of the molecule, a GraphCNN [13] based approach is suggested. The graph convolution layers used as a part of the “Molecule Network” (Figure 2) generate fixed length vector representations of the protein and ligand molecules. These will be provided as input to the reinforcement learning algorithm. The GraphCNN representation is expected to generate the molecular fingerprint by utilizing the atomic and spatial features of the protein ligand complex represented as graph node embedding.

**Fig. 2.**
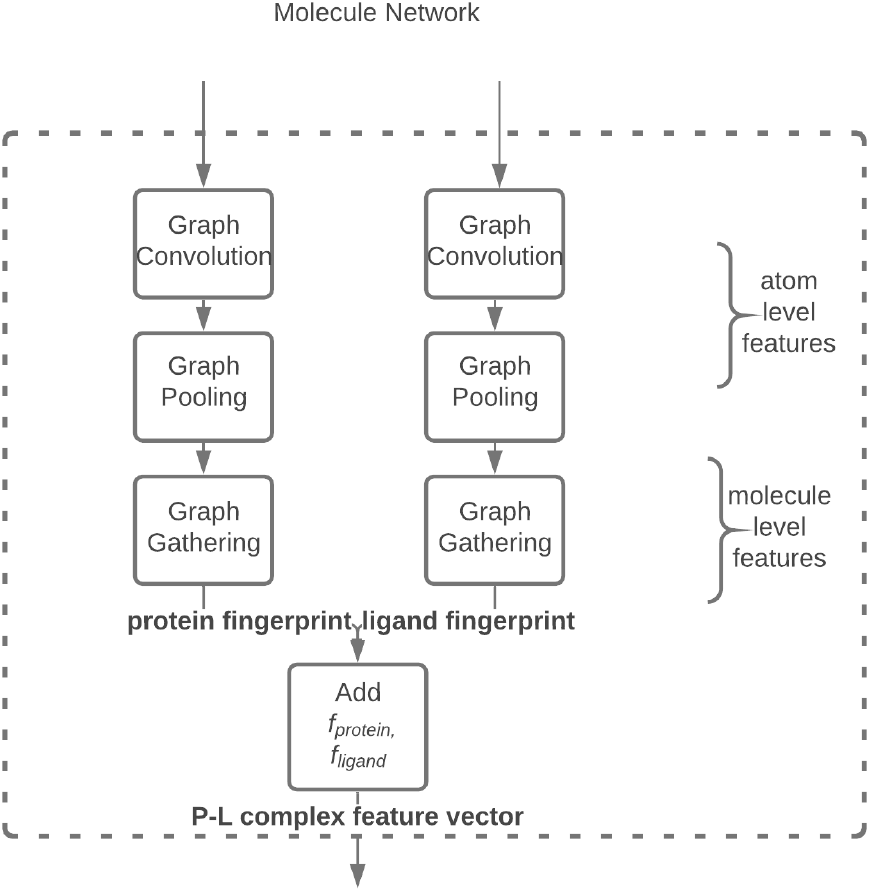
Molecule Representation: Input– Protein and Ligand Molecule representation, Output– Protein-Ligand Complex feature vector.

The fingerprint, represented as a feature vector, will be provided as the input to the reinforcement learning algorithm. The reinforcement learning agent will then learn which features from the fingerprint representation would generate a pose leading to an optimal docking. It is to be noted that the fingerprint generated is learned and the convolution layers will be trained as a part of the overall reinforcement learning training process.

#### 2) Deep Deterministic Policy Gradient

DDPG is a modelfree, off-policy reinforcement learning algorithm, which utilizes an actor-critic architecture. The actor function approximates optimal policy by mapping states to a specific action deterministically. The critic network is a neural network function approximator which utilizes the Bellman equations to learn the Q values of the input state and the corresponding actions, similar to the Q-learning algorithm. DDPG introduces a target network, which slowly tracks the learned actor-critic networks. The target network overcomes the convergence issue pertaining to the utilization of the same Q-network for policy update and target calculation.

#### 3) Optimization Policy

The DDPG algorithm tries to predict an optimal policy, which in the case of protein-ligand docking would be the optimal approximation of the docking function. The overall policy optimization function, represented as a maximization process for the reward ℝ, is denoted as:

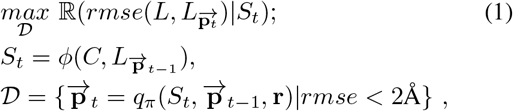

where the input *S* is a vector encoding of the atomic and molecular features generated from the protein *C* and the ligand *L* by the GraphCNN function *ϕ*, and 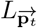 is the ligand with pose 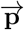 generated at time *t*, through a set of transitions 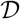 using the policy *q_π_*, which is the *state-value function* approximating the docking function, for a given stopping condition *rmse* < 2Å.

The optimization function in Eq (1) predicts the optimal ligand pose 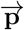 in the protein-ligand complex with the help of the atomic and molecular features represented as the feature vector *S*. Various molecular interactions including the active site interaction become an implicit property of the optimization function. This leads to a learned policy capable of predicting ligand poses within the protein’s active-site, thus incorporating the active-site prediction as an emergent optimization feature.

#### 4) Reward function

The goal of the reinforcement learning mechanism is to approximate an optimization function, such that the overall reward is maximized. In the case of proteinligand docking, the reinforcement learning algorithm approximates a pose prediction function such that the overall *rmse* is minimized. The reward function, thus, is a function of the *rmse* such that it behaves inversely to it, and maximizing the reward leads to a minimum *rmse* value.

Different reward functions have been suggested in the literature for reinforcement learning, each with its own benefits and shortcomings [34]. Some reward function forms that would be appropriate for this problem are shown in fig 3, and are described below:

- Inverse squared: This is the simplest form of the reward function, where ℝ(*x*) = 1/(*x*)^2^. The gradient of the reward function is very flat far from the goal, with a steep gradient as the agent nears the goal. This steep gradient helps the agent to quickly approach the goal, but the successive improvements in the policy are limited, as a small improvement in policy translates to a comparatively higher reward, thus slowing the progress once the agent has approached the goal.
- Sine hyperbolic: This reward function has the form ℝ(*x*) = 1/ sinh(*x^a^*) and behaves somewhat similarly to the inverse, however, the parameter *a* allows some control over the steepness of slope away from the maximum.
- Gaussian: This reward system is Gaussian inspired and has the form ℝ(*x*) = exp (− *x*^2^/(2*σ*^2^)). The reward function is uniformly distributed with a steep gradient when approaching the goal, and flattens out at the goal and away from it. This helps the agent to quickly approach the goal. The drawback of this reward function is that once the agent has approached the goal, it would prefer to revolve around the goal, rather than converging onto it, as the cumulative reward is higher near the goal as compared to *at* the goal due to the gradient. The parameter *σ* may be used to change the slope.
- Combination: It is possible to envisage a reward function created by combining the exponential and the sinh functions, so that its slope is steep both away from the maximum and close to the maximum. It is also possible to add a step function so that going far away from the maximum incurs not just a lower reward, but also a penalty. This would enable the RL algorithm to learn to stay near the maximum and also lead to faster optimization.
- Stochastic weight averaging: Averaging the reward function at each step over several preceding steps can reduce the learning instabilities caused by noisy gradient updates [35]. This also helps the RL to remember the highly rewarded policies, and to optimize accordingly.

**Fig. 3.**
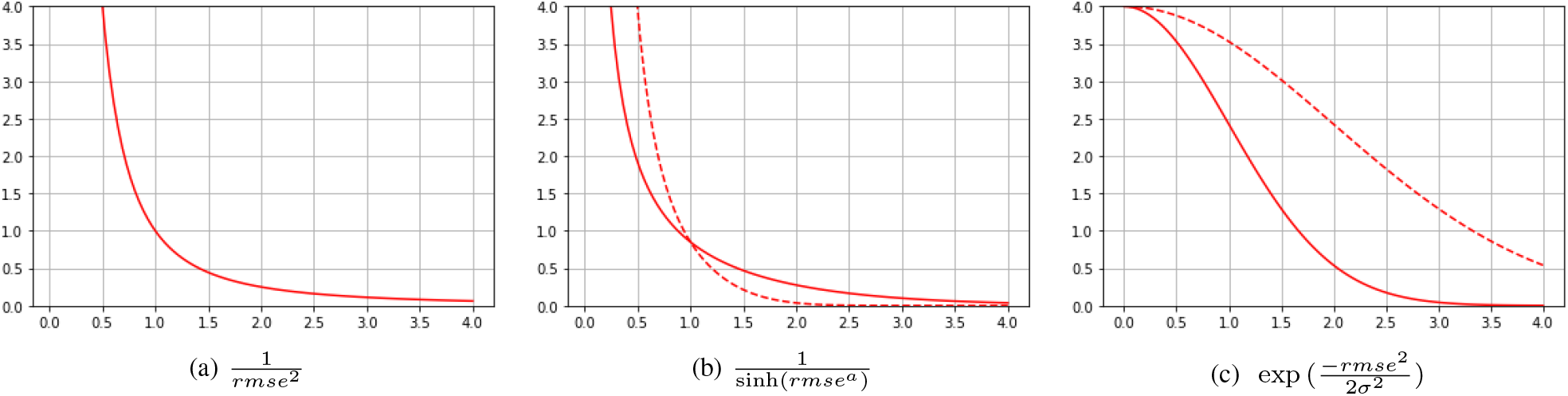
Reward as a function of the rmse

For an initial implementation of the DDPG algorithm, the Sine hyperbolic reward function may be chosen, since it has the attractive feature of a parameter that controls the steepness of its slope, and thus can be tuned to reach the goal quickly. Other reward functions can be tried based on the performance of this reward function.

#### 5) Training and Testing

The RL training is executed as an iterative process, which helps in generating scenarios in which the underlying policy network can be trained. As represented in figure 4, the training is done with known protein-ligand complexes. The input protein-ligand complex is separated out into protein and ligand representations. These separate molecular representations are used by the RL network to train the molecular graph and the corresponding optimal policy. In each of the training iterations, the RL network produces a ligand pose, which is compared against the original ligand in the protein-ligand complex for the *rmse* calculation. The *rmse* of the generated pose is compared to the terminating condition, which in this case is set to 2Å. In the case of an unfavourable *rmse*, ie. *rmse* > 2Å, the RL network gets updated as per the DDPG policy update equation *J*(*π*) = *E*(*r^γ^|π*), using a function of *rmse* as the policy reward. The training process would eventually produce an optimal policy network which generates an optimized ligand pose based on the features of the protein and ligand. The generated ligand pose is expected to be as close as possible to the original ligand pose in the protein-ligand complex due to the minimization of *rmse*.

**Fig. 4.**
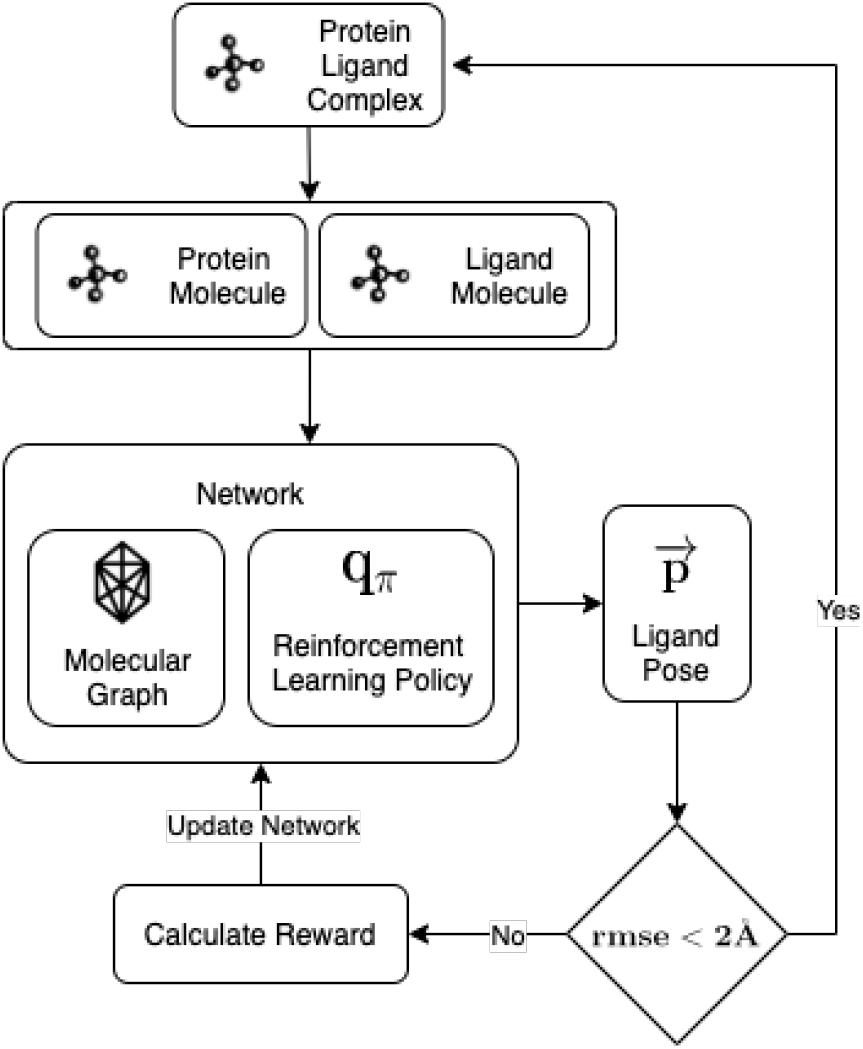
Reinforcement Learning Training Process: Input Protein Ligand complex, Output Reinforcement Learning Network.

The testing (fig. 5) also proceeds in an iterative manner. The RL Network in this case accepts a protein and ligand as input. Unlike the training phase, where the optimal ligand-pose is known, in testing phase, the optimal ligand pose is to be predicted; hence, *rmse* can no longer be used as a stopping condition for optimal ligand pose generation. Instead, 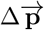 change in pose– becomes the deciding factor to ascertain if any better pose can be generated. Threshold Θ denotes a sufficiently low change in pose, 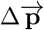, at which the generated pose is accepted as the optimal ligand pose.

**Fig. 5.**
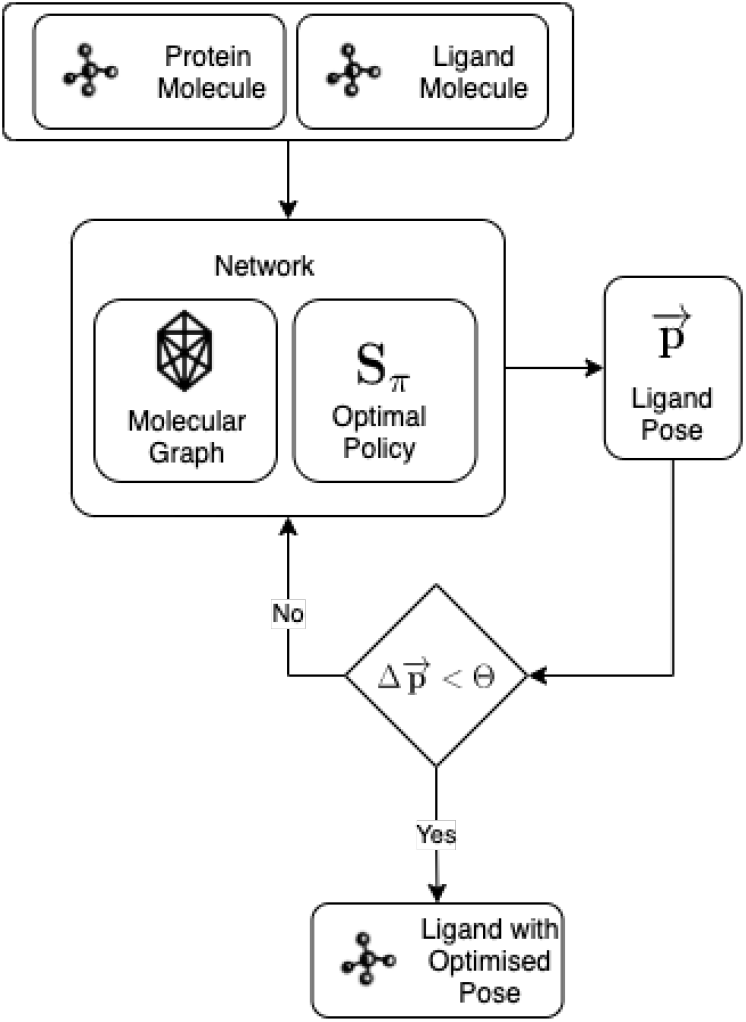
Reinforcement Learning Testing Process: Input Protein and Ligand molecules, Output Ligand with optimized pose with respect to Protein.

To validate the correctness of the generated optimal ligand-pose, an *rmse* comparison between the ligand pose of existing protein ligand complexes and the optimized ligand-pose as suggested by the RL model can be done.

Figure 6 represents the overall architectural design of the training network. The output of the training network is a target-actor network, which takes the protein and the ligand as inputs, and generates an optimal ligand pose as the output.

**Fig. 6.**
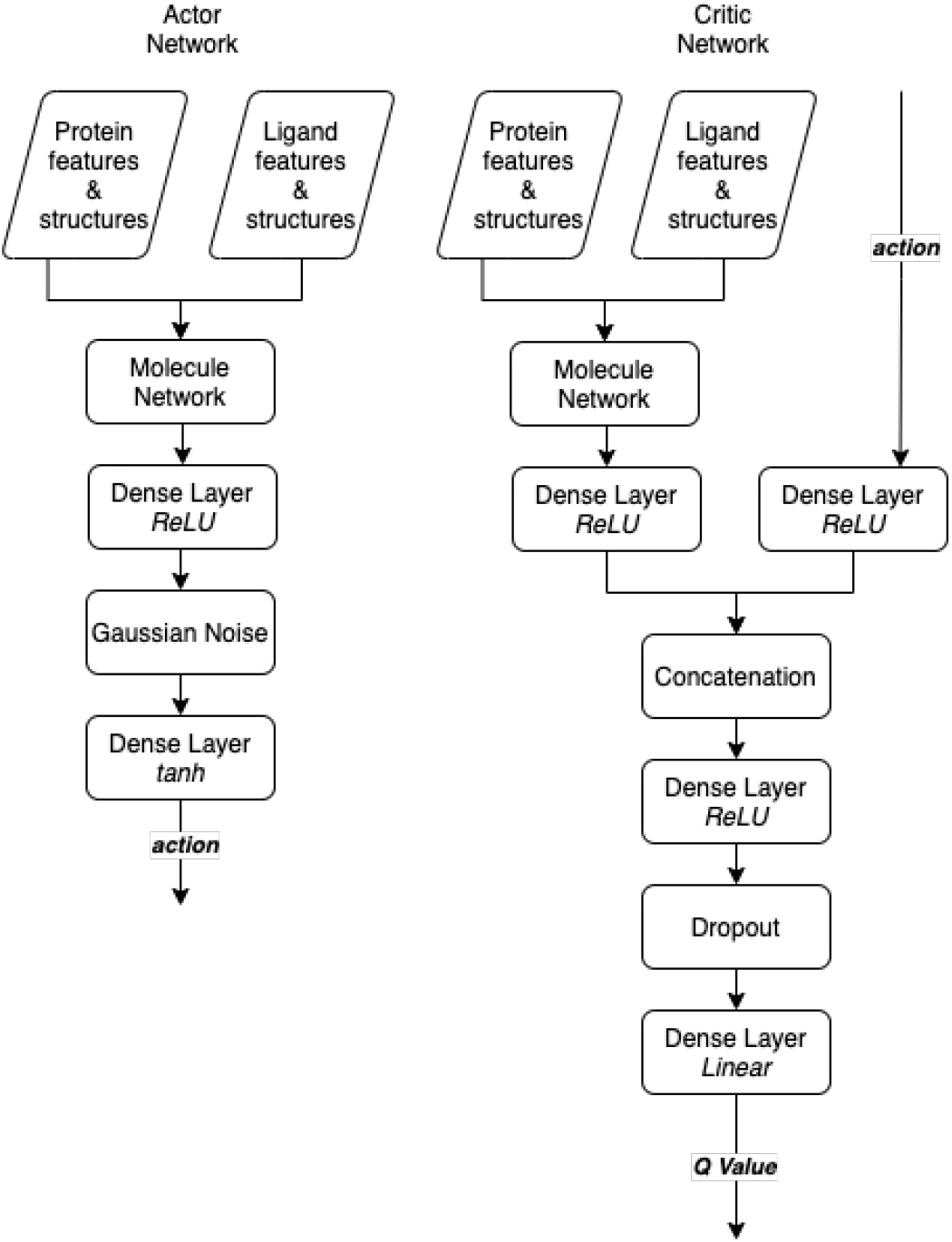
DDPG Network for pose prediction: Actor Network: Policy Network, Critic Network: Q-value network.

## IV. Discussions & Future Steps

The identification of correct binding pose in protein ligand interaction is of immense importance. The right binding pose is an indicator of true interactions between the amino acid residues and ligand molecule which eventually means the identification of required binding energy and binding affinities. The current state of the art methods like molecular docking utilize binding energy or binding affinity for finding the ligand pose. Often the predicted pose is not the biologically relevant pose, which could be due to various reasons. A known reason is that due to the compute power required to cover the entire chemical space for identifying the right ligand and ligand-pose, the biologically relevant pose may be missed by the docking algorithm. The RL approach suggested in this paper mathematically approximates a function to generate ligand docking poses utilising the protein and ligand atomic features, which would be learned from existing protein-ligand complexes. This is expected to reduce the overall search space for the identification of the right docking pose, hence making the docking process computationally cheaper compared to traditional methods. This method also eliminates the need to predefine the active sites in the protein, since the active site is predicted as part of the optimization process in this formalism.

The RL approach discussed in this paper brings in a new challenge with respect to molecule representation. The 3D CNNs provide a very elegant way of preserving the structural representation of the protein and ligand in spatial domain, but at a computational efficiency trade-off. This trade-off can be mitigated by using a graphical representation with the help of GraphCNN, which has shown great promise in terms of computational effectiveness in comparison to the 3D convolution network. The GraphCNNs are also good for representing sparse connected data. The GraphCNN representation preserves the atomic interactions through connected graph edges, but loses out on the structural representation. In our suggested approach, we mitigate this by encoding the spatial features as node features. This approach of combining node features with spatial features has not yet been attempted in the existing literature. If successful, this can certainly present a novel approach to molecular representation for protein-ligand interactions.

We can envision certain challenges with the proposed approach, primarily due to the data representation and the underlying machine learning algorithms used. In the GraphCNN representation, the 3D molecular structure becomes latent in the 2D representation, thus obscuring the learning of the convolutional section of the network. Also, our molecular representation groups together spatial and atomic features, a method that has not been previously attempted, hence its success needs to be validated. We choose DDPG as our RL algorithm, since the state-space is continuous. However, DDPG has been known to suffer from convergence issues [36]. The algorithm could converge sub-optimally or even diverge from solution if rewarding policies are not identified early on in training, making proper selection of reward function vital to its success. Even in the absence of reward, actor and critic training may trigger non-negligible updates that cause the actor-critic network to saturate very quickly. Hyperparameter tuning, choice of initial weights, and especially a stable noise model, are some requirements to prevent early saturation.

An alternate approach to the problem could be to discretize the continuous state space. By doing this, we open up the possibility of using other discrete-space RL models such as Double Deep Q Networks (DDQN). One may also use a more sophisticated continuous space algorithm, such as the Twin Delayed DDPG (TD3) [36]. These can be valid options to explore during implementation.

In the implementation stage of this problem, we expect to train a model based on variety of viral protein-ligand complexes. At this stage, we would also need to validate the supposition that the RL algorithm would be able to successfully identify the correct active site in the protein, and evaluate the importance of the various atomic features in the training process. It is also possible to explore ways to encode molecular features, e.g., features specific to ligands and residues, into the algorithm.

In summation: even though there are plenty of algorithms present for protein ligand interactions, identification of the right ligand pose is still dependent on various manual tasks. We propose a reinforcement learning based method where the agent optimizes the right pose and can also be trained to find the binding site. This will eliminate the need of identification of protein pocket and grid to position the ligand. The initial implementation of this formalism will be focused on testing the RL agent’s capabilities for antiviral protein ligand complexes. Eventually this can be extended to a more dynamic framework which is trained on multiple protein ligand data sets for predicting the optimized pose while identifying the binding site.

## Availability

The code for running smaller experiments for reinforcement learning based approach to protein ligand interaction and the dataset used is available at https://gitlab.com/lifesciences/rl-virtual-screening

## Acknowledgements

We would like to thank Dr. Manali Joshi, Dr. B. Jayaram and Ms. Shruti Koulgi for great discussions and suggestions. We would like to acknowledge Drug Discovery Hackathon organizers and Dr. Karthik Raman for giving us this opportunity to participate in one of the cutting edge problem statement and being approachable for our queries regarding the problem statement.

## Footnotes

1 https://innovateindia.mygov.in/ddh2020/

